# Individual differences in visual salience vary along semantic dimensions

**DOI:** 10.1101/444257

**Authors:** Benjamin de Haas, Alexios L. Iakovidis, D. Samuel Schwarzkopf, Karl R. Gegenfurtner

## Abstract

What determines where we look? Theories of attentional guidance hold that image features and task demands govern fixation behaviour, while differences between observers are ‘noise’. Here, we investigated the fixations of > 100 human adults freely viewing a large set of complex scenes. We found systematic individual differences in fixation frequencies along six semantic stimulus dimensions. These differences were large (> twofold) and highly stable across images and time. Surprisingly, they also held for *first* fixations directed towards each image, commonly interpreted as ‘bottom-up’ visual salience. Their perceptual relevance was documented by a correlation between individual face salience and recognition skills. The dimensions of individual salience and their covariance pattern replicated across samples from three different countries, suggesting they reflect fundamental biological mechanisms of attention. Our findings show stable individual salience differences along semantic dimensions, with meaningful perceptual implications. Salience reflects features of the observer as well as the image.

## Introduction

Humans constantly move their eyes^1^. The foveated nature of the human visual system balances detailed representations with a large field of view. On the retina^2^ and in visual cortex^3^, resources are heavily concentrated towards the central visual field, resulting in the inability to resolve peripheral clutter^4^ and the need to fixate visual objects of interest. Where we move our eyes determines which objects and details we make out in a scene^5,6^.

Models of attentional guidance aim to predict which parts of an image will attract fixations based on image features^7–10^ and task demands^11,12^. Classic salience models compute image discontinuities of low level attributes, such as luminance, colour and orientation^13^. These low-level models are inspired by ‘early’ visual neurons and their output correlates with neural responses in subcortical^14^ and cortical^15^ areas thought to represent neural ‘salience maps’. However, while these models work relatively well for impoverished stimuli, human gaze behaviour towards richer scenes can be predicted at least as well by the locations of objects^16^ and perceived meaning^9^. When sematic object properties are taken into account, their weight for gaze prediction far exceeds that of low-level attributes^8,17^. A common thread of low- and high-level salience models is that they interpret salience as a property of the image and treat inter-individual differences as unpredictable^7,18^, often using them as a ‘noise ceiling’ for model evaluations^18^.

However, even the earliest studies of fixation behaviour noted considerable individual differences^19,20^ and basic occulomotor traits, like mean saccadic amplitude and velocity, reliably vary between observers^21–27^. Moreover, recent twin-studies revealed that social attention and gaze traces across complex scenes are highly heritable^28,29^. This suggests individual differences in fixation behaviour are not random, but systematic. However, it is largely unclear, *how* individuals differ in their fixation behaviour and what may explain these differences. Can individual fixation behaviour be captured along a limited set of dimensions?

Here, we tested the hypothesis that individual gaze reflects individual *salience differences* along a limited number of semantic dimensions. We investigated the fixation behaviour of > 100 human adults freely viewing 700 complex scenes, containing thousands of semantically annotated objects^8^. We quantified salience differences as the *individual* proportion of cumulative fixation time or *first* fixations landing on objects with a given semantic attribute. In free viewing, the *first* fixations after image onset are thought to reflect ‘automatic’ or ‘bottom-up’ salience^30–32^, especially for short saccadic latencies^33,34^. They may therefore reveal individual differences with a deep biological root. We tested the reliability of such differences across random subsets of images and re-tests after several weeks. To test the generalizability of salience differences, we replicated their set and covariance pattern across independent samples from three different countries. Finally, we explored whether individual salience differences are related to personality and perception, focussing on the example of face salience and face recognition skills for the latter.

## Results

### Reliable Salience Differences along Semantic Dimensions

We tracked the gaze of healthy human adults freely viewing a broad range of images depicting complex everyday scenes^8^. A first sample was tested at University College London, UK (*Lon*; *n* = 51), and a replication sample at University of Giessen, Germany (*Gi_1*; n = 51). The replication sample was also invited for a re-test after two weeks (*Gi_2*; *n* = 48). Additionally we re-analysed a public dataset from Singapore (*Xu et al*. ^8^; *n* = 15).

First, we probed the individual tendency to fixate objects with a given semantic attribute, measuring duration-weighted fixations across a free-viewing period of 3s. We considered a total of twelve semantic properties, which have previously been shown to carry more weight for predicting gaze behaviour (on an aggregate group level) than geometric or pixel-level attributes^8^. To test the consistency of individual salience differences across independent sets of images, we probed their reliability across 1000 random (half-)splits of 700 images. We found consistent individual salience differences (*r* > .6) for six of the 12 semantic attributes: Neutral *Faces*, *Emotion*al Faces, *Text*, objects being *Touched*, objects with a characteristic *Taste* (i.e. food and beverages) and objects with implied *Motion* (Figure 1, grey scatter plots).

**Figure 1.**
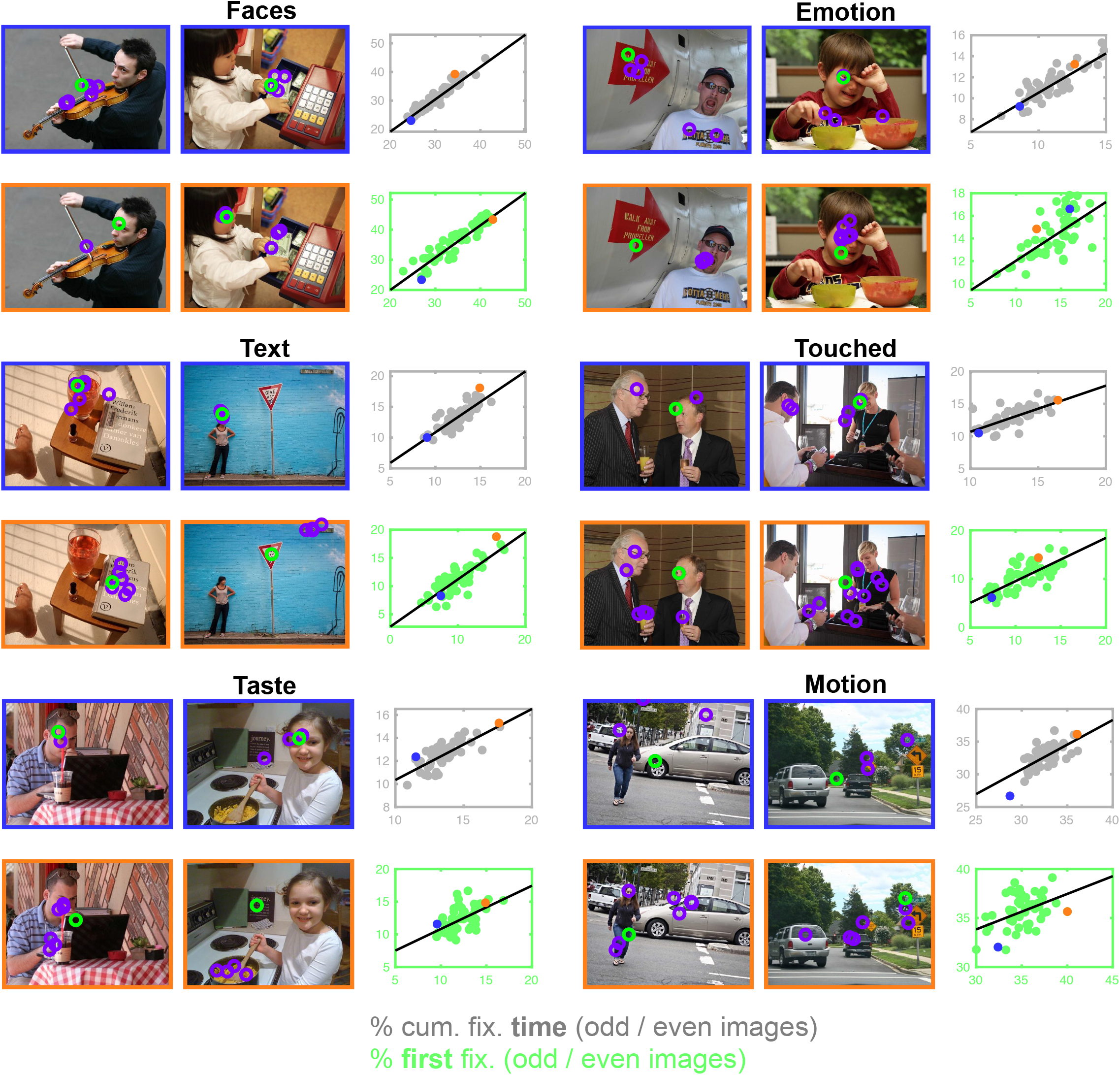
Consistent individual differences in fixation behaviour along six semantic dimensions. For each semantic attribute, the grey scatter plot shows individual proportions of cumulative fixation time for the odd *versus* even numbered images in the *Lon* dataset. The green scatter plot shows the corresponding individual proportions of *first* fixations after image onset. For each dimension, two example images are given and overlaid with the fixations from one observer strongly attracted by the corresponding attribute (orange frames) and one observer weakly attracted by it (blue frames). The overlays show the first fixation after image onset as a green circle; any subsequent fixations are shown in purple. The two data points corresponding to the example observers are highlighted in the scatter plot, corresponding to the colour of the respective image frames.

Observers showed up to two-fold differences in the cumulative fixation time attracted by a given semantic attribute and the median consistency of individual differences across image splits for these six dimensions ranged from *r* = .64 *P* < .001 (*Motion*) to r = .94, *P* < .001 (*Faces*; Table 1, left hand side; *P*-Values Bonferroni corrected for 12 consistency correlations).

**Table 1.** Range and median consistency of individual differences in fixation behaviour towards six semantic dimensions. The left hand side of the table shows data for individual differences in the proportion of cumulative fixation time (across three seconds) spent on objects with the respective semantic attributes. The right hand side shows data on individual differences in the proportion of first fixations after image onset landing on objects with those attributes. For each semantic attribute, the data from different samples (*Lon*, *Gi_1, Gi_2* and *Xu et al*.) is presented in separate rows. The range across participants is given in % fixation time or % of first fixations and the max/min ratio indicates the relative difference between observers attracted the most or least by a given attribute. Pearson correlations (*r*) indicate the median split-half correlation of individual differences across 1000 random image splits and the corresponding *P*-value (Bonferroni corrected for 12 semantic attributes in the *Lon* sample, as indicated by the *). Consistency correlations failing to reach statistical significance are highlighted in red. The final row for each attribute shows the corresponding re-test reliability across several weeks.

|  | Ind. % fixation time |  |  |  | Ind. % first fix |  |  |  |
| --- | --- | --- | --- | --- | --- | --- | --- | --- |
|  | *r* | *P* | range [%] | max/min | *r* | *P* | range [%] | max/min |
| **Faces - Lon** | .94 | <.001* | 24-43 | 1.80 | .88 | <.001* | 24-43 | 1.77 |
| – Gi_1 | .91 | <.001 | 21-35 | 1.64 | .88 | <.001 | 15-42 | 2.82 |
| – Gi_2 | .91 | <.001 | 17-34 | 1.99 | .89 | <.001 | 15-43 | 2.76 |
| – Xu *et al.* | .95 | <.001 | 23-35 | 1.50 | .86 | <.001 | 24-41 | 1.69 |
| - Re-test (Gi_1-2) | .80 | <.001 | - | - | .86 | <.001 | - | - |
| **Emotion - Lon** | .82 | <.001* | 8-15 | 1.93 | .69 | <.001* | 9-18 | 1.88 |
| – Gi_1 | .80 | <.001 | 8-13 | 1.68 | .76 | <.001 | 8-17 | 2.20 |
| – Gi_2 | .82 | <.001 | 7-13 | 1.79 | .75 | <.001 | 8-17 | 2.22 |
| - Xu *et al.* | .89 | <.001 | 8-12 | 1.56 | .48 | .07 | 13-17 | 1.31 |
| - Re-test (Gi_1-2) | .85 | <.001 | - | - | .79 | <.001 | - | - |
| **Text - Lon** | .81 | <.001* | 9-16 | 1.76 | .77 | <.001* | 6-17 | 2.96 |
| – Gi_1 | .90 | <.001 | 9-19 | 1.97 | .82 | <.001 | 5-16 | 3.11 |
| – Gi_2 | .91 | <.001 | 9-20 | 2.21 | .82 | <.001 | 4-17 | 3.91 |
| - Xu *et al.* | .94 | <.001 | 8-18 | 2.16 | .83 | <.001 | 4-13 | 3.25 |
| - Re-test (Gi_1-2) | .84 | <.001 | - | - | .89 | <.001 | - | - |
| **Touched - Lon** | .73 | <.001* | 11-16 | 1.52 | .73 | <.001* | 6-16 | 2.48 |
| - Gi_1 | .74 | <.001 | 9-15 | 1.63 | .71 | <.001 | 6-15 | 2.65 |
| - Gi_2 | .75 | <.001 | 9-16 | 1.67 | .72 | <.001 | 6-16 | 2.74 |
| - Xu *et al.* | .91 | <.001 | 9-16 | 1.85 | .65 | .008 | 7-13 | 1.81 |
| - Re-test (Gi_1-2) | .70 | <.001 | - | - | .78 | <.001 | - | - |
| **Taste - Lon** | .70 | <.001* | 10-16 | 1.58 | .57 | <.001* | 10-16 | 1.67 |
| - Gi_1 | .70 | <.001 | 11-15 | 1.38 | .29 | .04 | 10-14 | 1.36 |
| - Gi_2 | .74 | <.001 | 10-17 | 1.67 | .50 | <.001 | 9-16 | 1.75 |
| - Xu *et al.* | .84 | <.001 | 11-16 | 1.44 | .69 | .004 | 9-15 | 1.72 |
| - Re-test (Gi_1-2) | .78 | <.001 | - | - | .62 | <.001 | - | - |
| **Motion - Lon** | .64 | <.001* | 28-36 | 1.31 | .35 | .15* | 31-39 | 1.25 |
| - Gi_1 | .65 | <.001 | 28-34 | 1.22 | .52 | <.001 | 30-38 | 1.29 |
| - Gi_2 | .73 | <.001 | 26-33 | 1.29 | .52 | <.001 | 28-37 | 1.32 |
| - Xu *et al.* | .85 | <.001 | 28-33 | 1.19 | .62 | .01 | 33-40 | 1.22 |
| - Re-test (Gi_1-2) | .68 | <.001 | - | - | .78 | <.001 | - | - |

Previous studies have argued that extended viewing behaviour is governed by cognitive factors, while first fixations towards a free-viewed image are governed by ‘bottom-up’ salience^30–32^, especially for short saccadic latencies^33,34^. Others have found that perceived meaning^9,35^ and semantic stimulus properties^8,36^ are important predictors of gaze behaviour from the first fixation. We found consistent individual differences also in the proportion of *first* fixations directed towards each attribute. The range of individual differences in the proportion of *first* fixations directed to each of the six attributes was even larger than that for all fixations (up to threefold). Importantly, these inter-observer differences were consistent for all dimensions found for cumulative fixation time except *Motion* (*r* = .34, *n.s*.), ranging from *r* = .57, *P* < .001 (*Taste*) to r = .88, *P* < .001 (*Faces*; Table 1, right hand side and green scatter plots in Figure 1; *P*-Values Bonferroni corrected for 12 consistency correlations).

These salience differences proved robust for different splits of images and replicated across datasets from three different countries (Figure 2a; Table 1). For the confirmatory *Gi_1* dataset we tested the same number of observers as in the original *Lon* set. A power analysis confirmed that this sample size yields > 95% power to detect consistencies with a population effect size of *r* > .5. For cumulative fixation time (grey histograms in Figure 2a), the six dimensions identified in the *Lon* sample (top row), closely replicated in the *Gi_1* and *Gi_2* samples (middle rows), as well as in a re-analysis of the public *Xu et al*. dataset (bottom row), with consistency correlations ranging from .65 (*Motion* in the *Gi_1* set) to .95 (*Faces* in the *Xu et al*. set; left column of Table 1). Similar was true for *first* fixations, although the consistency correlation for *Emotion* missed statistical significance in the small *Xu et al*. dataset (green histograms in Figure 2a; right column of Table 1).

**Figure 2.**
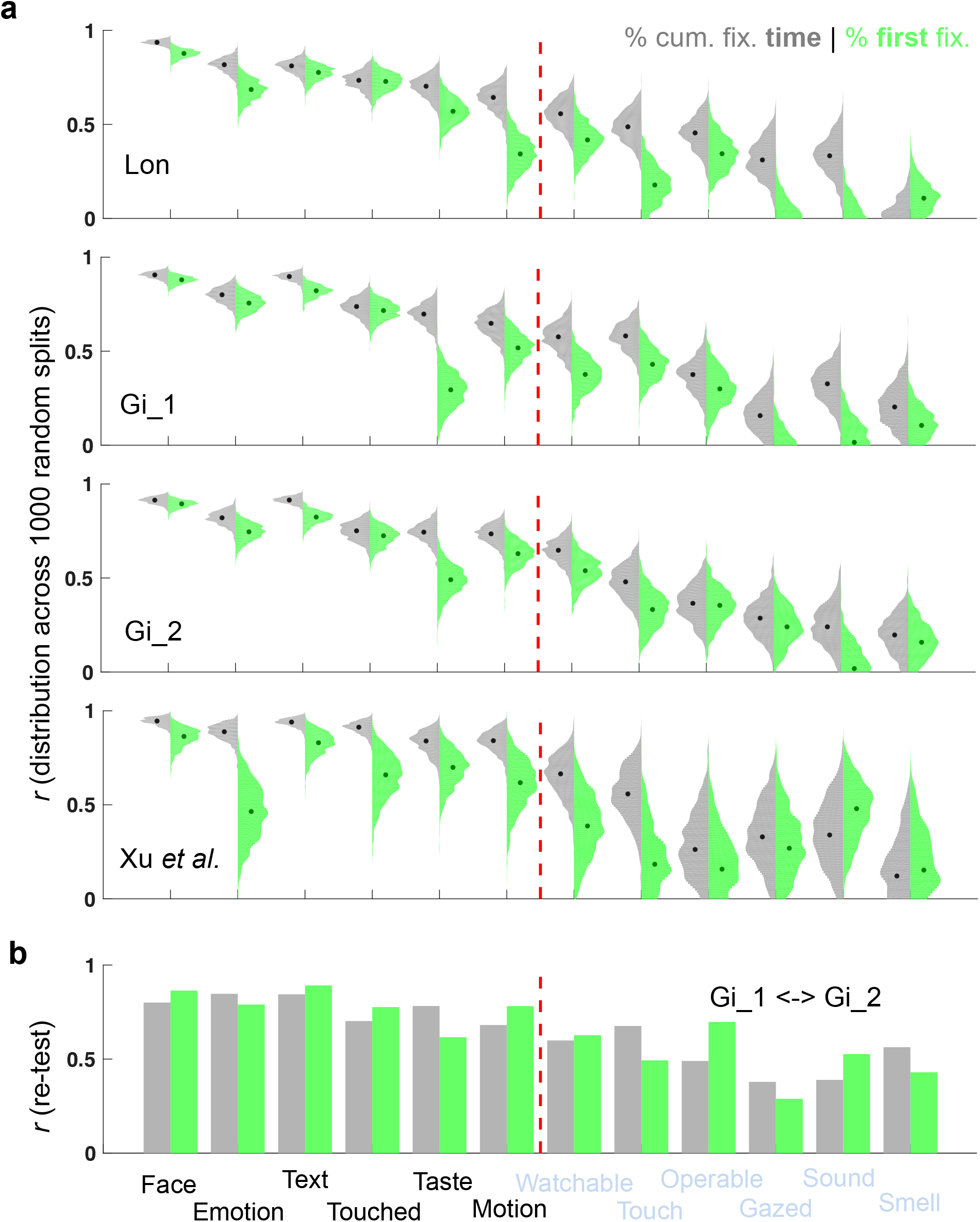
Consistency of results across images, datasets and time. **(a)** Distribution plots of consistency correlations for each of the twelve semantic dimensions tested (as indicated by the labels on the x-axis in b). Results are shown separately for the four observer samples tested (as indicated by the row labels). Gi_1 and Gi_2 refer to the first and second appointment of the Gi sample. The grey left-hand leaf of each distribution plot shows a histogram of split-half correlations for 1000 random splits of the image set, the green right hand leaf shows the corresponding histogram for *first* fixations after image onset. Overlaid dots indicate the median consistency correlation for each distribution. High split-half correlations indicate consistent individual differences in fixation behaviour across images for a given dimension. The dashed red line separates the six attributes found to be consistent dimensions of individual differences in the *Lon* sample from the remaining dimensions. (**b**) Re-test reliability across Gi_1 and Gi_2. The magnitude of re-test correlations for individual dwell time and proportion of first fixations is indicated by grey and green bars, respectively. All correlation and *P*-values can be found in Table 1.

The individual salience differences we found were consistent across subsets of diverse, complex images. To test whether they reflected stable observer traits, we additionally tested their re-test reliability for the full image set across a period of 6-43 days (average 16 days; the *Gi_1* and *Gi_2* datasets). Salience differences along all six semantic dimensions were highly consistent over time (Figure 2b). This was true for both, cumulative fixation time (re-test reliabilities ranging from *r* = .68, *P* < .001 (*Motion*) to r = .85, *P* < .001 (*Faces*); left column of Table 1; grey bars in Figure 2b) and *first* fixations (re-test reliabilities ranging from *r* = .62, *P* < .001 (*Taste*) to r = .89, *P* < .001 (*Text*); right column of Table 1; green bars in Figure 2b).

Finally, we tested the relationship between individual salience and visual field biases. Different types of objects tend to appear at different parts of the visual field and recent findings suggest that observers are attuned to these contingencies^37–39^. The most prominent spatial bias in the scene stimuli we used was a strong tendency for faces to appear in the upper visual field (~ 75%). Individual face salience (% *first* fixations) was indeed correlated with a general upper visual field bias (as indexed by the median elevation of *first* fixations towards images not containing a face; *r* = .49-.78 across samples, all *P* < .05). Crucially, however, individual differences in face salience persisted independent of this spatial bias. Individual face salience in the *lower* and upper visual field were highly correlated with each other (*r* = .61-.87 across samples, all *P* < .05; see Supplementary Methods and Supplementary Fig. 1 for details).

### Covariance Structure of Individual Differences in Semantic Salience

Having established reliable individual differences in fixation behaviour along semantic dimensions, we further explored the space of these difference by quantifying the covariance between them. For this analysis we collapsed neutral and emotional faces into a single *Faces* label, because they are semantically related and corresponding differences were strongly correlated with each other (r = .74, *P* < .001; *r* = .81, *P* < .001 for cumulative fixation times and first fixations, respectively). Note that we decided to keep these two dimensions separated for the analyses above because the residuals of fixation times for emotional faces still varied consistently when controlling for neutral faces (r = .73, *P* < .001), indicating an independent component (however the same was not true for *first* fixations, r = .24, *n.s*.).

The resulting five dimensions showed a pattern of pairwise correlations that allowed the identification of two clusters (Fig. 3b, left hand side). This was illustrated by the projection of the pairwise (dis-)similarities onto a two-dimensional space, using metric dimensional scaling (Fig. 3b, right hand side). *Faces* and *Motion* were positively correlated with each other, but negatively with the remaining three attributes *Text*, *Touched* and *Taste*. Interestingly, *Faces*, the most prominent dimension of individual fixation behaviour, was strongly anti-correlated with *Text* and *Touched*, the second and third most prominent dimensions (*Text*: r = -.62, *P* < .001 and r = -.47, *P* < .001 for cumulative fixation times and first fixations, respectively, Fig. 3a, left hand side; *Touched*: r = -.58, *P* < .001 and r = -.80, *P* < .001, Fig. 3a, right hand side). These findings closely replicated across all four datasets (Fig. 4). Pair-wise correlations between (z-converted) correlation matrices from different samples ranged from .68 to .95 for cumulative fixation times and from .91 to .98 for first fixations.

**Figure 3.**
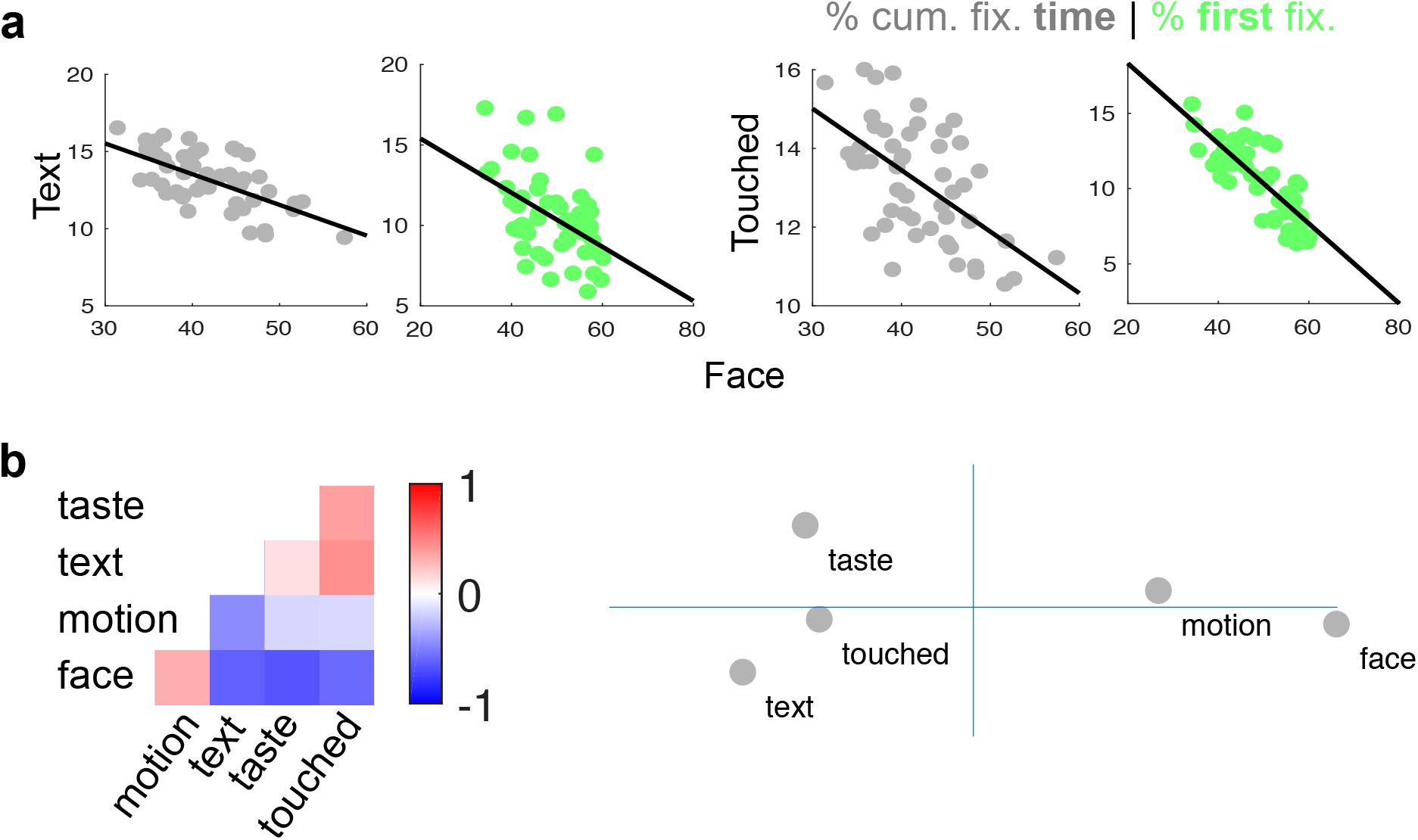
Covariance of individual differences along semantic dimensions. (**a**) Grey scatter plots show the individual proportion of cumulative fixation time (in %) for *Faces versus Text* (left hand side) and *Faces versus* objects being *Touched* (right hand side). Green scatter plots show the corresponding data for the individual proportion of *first* fixations after image onset. (**b**) Correlation matrix for individual differences along five semantic dimensions (left hand side; note that the labels for emotional and neutral faces were collapsed for this analysis). Colour indicates pairwise Pearson correlation coefficients as indicated by the bar. *Motion* and *Face* are positively correlated with each other, but negatively correlated with the remaining dimensions, as captured by a two-cluster solution of metric dimensional scaling to two dimensions (right hand side). All data shown are based on individual proportions of fixation time in the *Lon* dataset. The corresponding supplemental Figure S1, shows the consistency of this pattern for the proportion of first fixations and across all four datasets.

**Figure 4.**
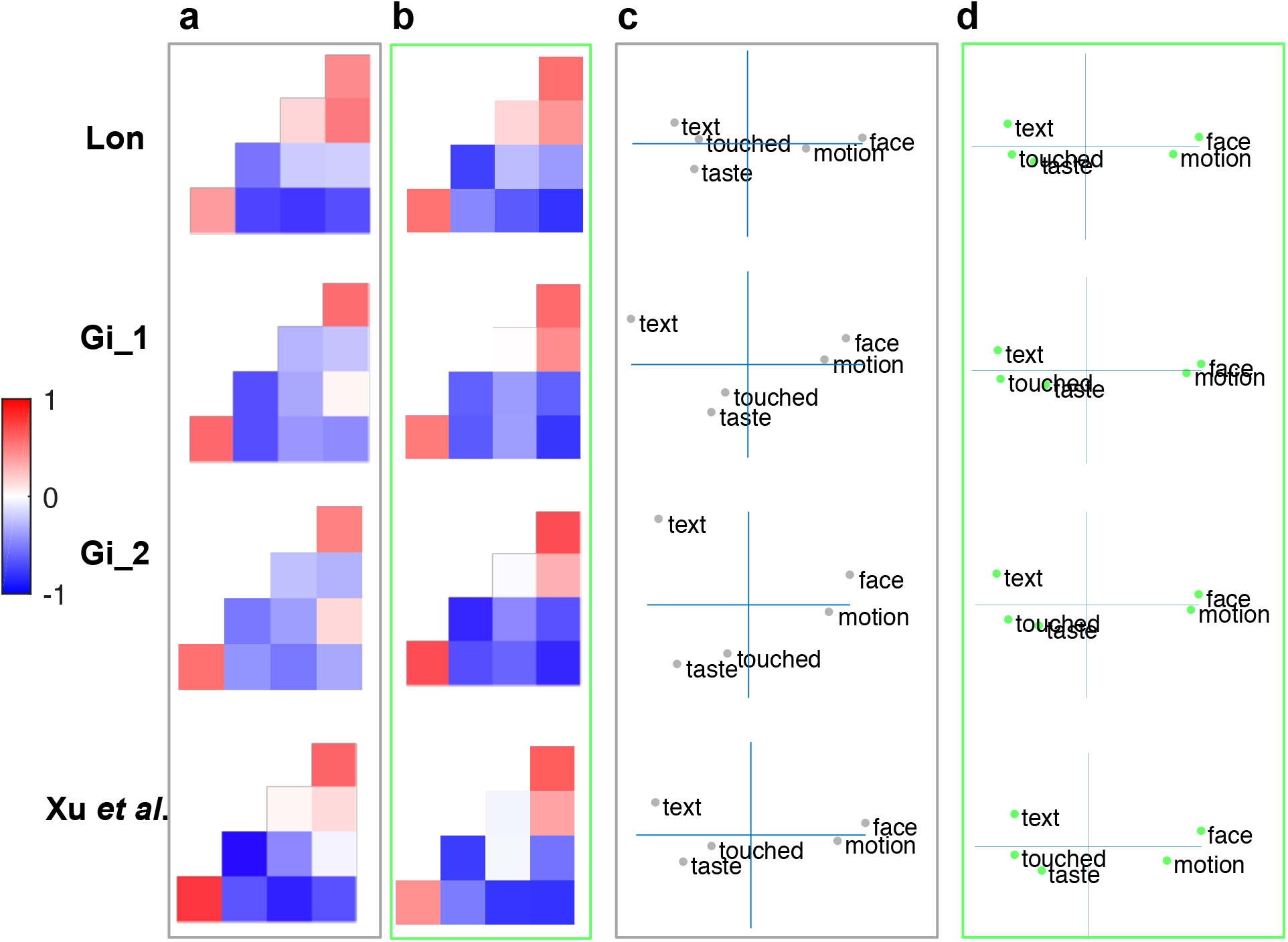
Consistency of covariance pattern across datasets. (**a**) shows covariance patterns for dimensions of individual differences in cumulative fixation durations and (**b**) those for individual differences in the proportion of*first* fixations. Each row shows the results for one dataset as indicated (Lon, Gi_1, Gi_2 and Xu *et al*.). Colours of the left hand correlation matrices indicate pairwise Pearson correlation coefficients between dimensions as indicated by the bar. Please refer to Figure 3 for dimension labels (from top to bottom: taste, text, motion, face; from left to right: motion, text, taste, touched). Note the similarity of covariance patterns between datasets. All pairwise correlations between (Fisher z-transformed) correlation matrices were >.68 for cumulative fixation durations and >.90 for first fixations. The right hand side scatter plots show the results of metric multidimensional scaling onto two dimensions for cumulative fixation duration (**c**) and first fixations (**d**). This analysis yielded a distinct cluster for *Faces* and *Motion*, as well as a cluster for *Text*, *Touched Taste* for first fixations in all datasets. This structure was very similar for cumulative dwell time, with the exception of *Text* separating from *Touched* and *Taste* in the Gi_1 and Gi_2 data.

### Perceptual and Personality Correlates of Salience Differences

If salience differences are indeed deeply rooted in the visual cortices of our observers, then this might have an effect on their perception of the world. We aimed to test this hypothesis by focussing on the most prominent dimension of salience differences: *Faces*, as indexed by the individual proportion of *first* fixations landing on faces (which is thought to be an indicator of bottom-up salience). 46 observers from the *Gi* sample took the Cambridge Face Memory Test (CFMT) and we tested the correlation between individual face salience and face recognition skills. CFMT scores and the individual proportion of *first* fixations landing on faces correlated with *r* = .41, *P* < .005. Interestingly, this correlation did not hold for the individual proportion of total cumulative fixation time landing on faces, which likely represents more voluntary differences in viewing behaviour (*r* = .21, *n.s*.; Fig. 5a).

**Figure 5.**
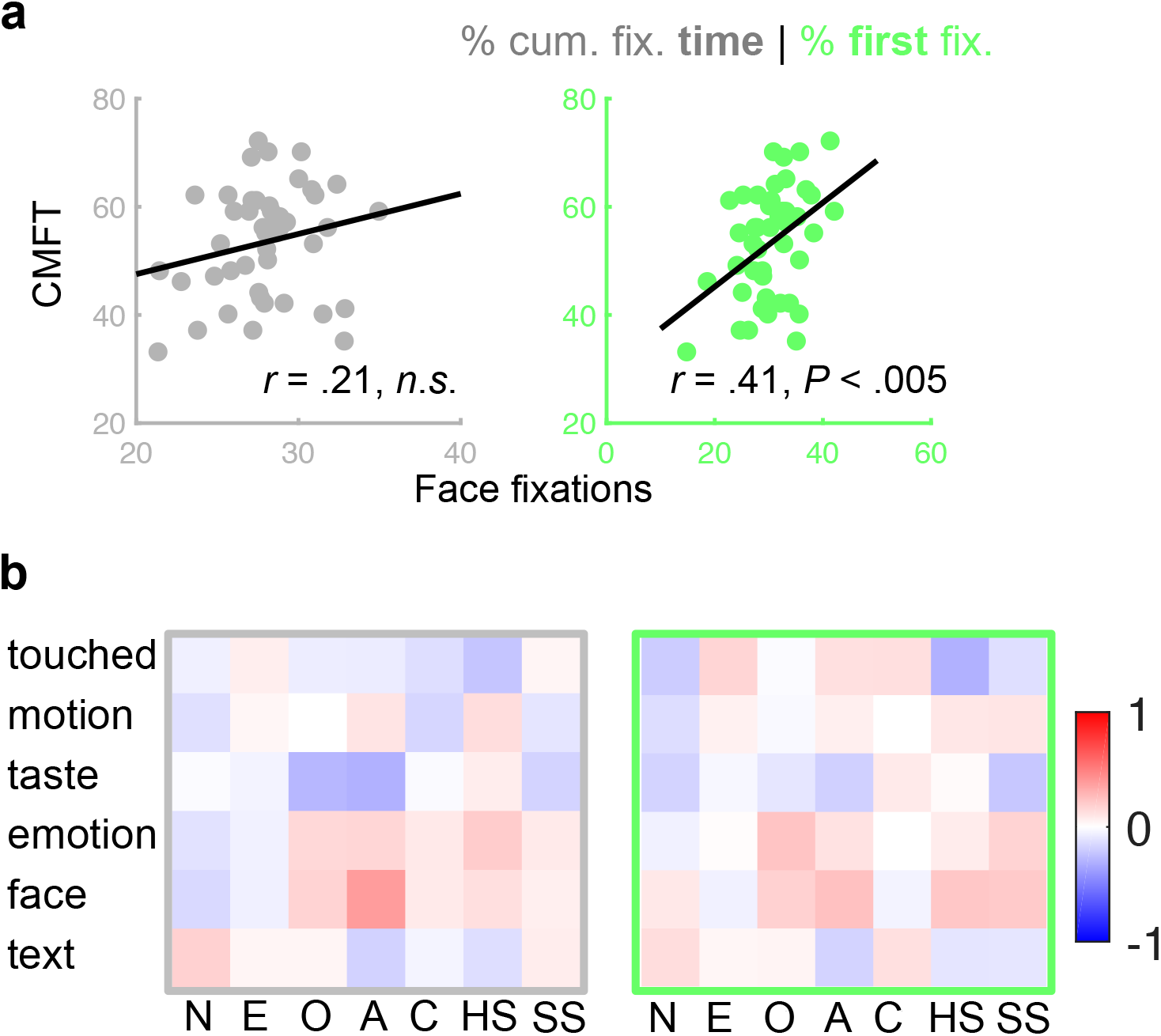
Correlations between fixation behaviour, face recognition skills and personality. (**a**) Performance on the Cambridge Face Memory Test (CMFT) correlated significantly with the individual proportion of first fixations landing on faces when freely viewing complex scenes (green, right hand side). This correlation was not significant for cumulative fixation time across the viewing time of three seconds (grey, left hand side). (**b**) Neither cumulative fixation time (grey frame, left hand side), nor the proportion of first fixations (green frame, right hand side) towards any of the six dimensions of individual gaze behaviour correlated significantly with any of the tested dimensions of personality (N: Neuroticism, E: Extraversion, O: Openness, A: Assertiveness, C: Conscientiousness, HS: High Sensitivity, SS: Sensation Seeking). All data from the Gi_1 sample.

Having established a correlation of semantic salience differences with perception, we then explored potential relationships with personality variables. Observers in the *Lon* sample completed questionnaires for seven personality dimensions, including the Big Five^40^, Sensation Seeking^41^ and High Sensitivity^42^ (see Methods for details). We tested all 7×6 pairwise correlations between personality traits and individual salience differences, applying family-wise error correction for multiple comparisons^43^. Results showed no significant relationship between personality variables and cumulative fixation times (all *r* < .39, *n.s*.; Fig. 5b, left hand side), or *first* fixations (all *r* < .23, *n.s*.; Fig. 5b, right hand side).

## Discussion

Individual differences in gaze traces have been documented since the earliest days of eyetracking^19,20^. However, salience models have routinely ignored them and the nature of these differences was unclear. Our findings show that what was thought to be noise can actually be explained by a canonical set of semantic salience differences. These salience differences were highly consistent across hundreds of complex scenes, proved reliable in a re-test after several weeks and persisted independently of correlated visual field biases. This shows that visual salience is not just a factor of the image; individual salience differences are a stable trait of the observer.

Not only the set of these differences, but also their covariance structure replicated across independent samples from three different countries. This may partly be driven by environmental and image statistics (for instance, faces are more likely to move than food). But it probably also reflects a deep-rooted neurobiological basis of these differences. This possibility is underscored by earlier studies showing that the visual salience of social stimuli is reduced in individuals with autism spectrum disorder^28,44,45^. Most importantly, recent twin studies in infants and children show that individual differences in gaze traces are heritable^28,29^. The gaze trace dissimilarities investigated in these twin studies might be a manifestation of the salience differenceswe found here. This suggests a strong genetic component for individual salience differences.

Recent findings in macaque show that face deprived monkeys have a drastically reduced fixation tendency towards faces and an increased fixation tendency for hands. These differences in fixation behaviour were accompanied by an underdevelopment of domain specific face patches in the temporal cortex and a relative overdevelopment of hand preferring patches^46^. In our study, high individual salience for the *Touched* label implies strong attentional guidance by hands (see Fig. 1 for examples). It is tempting to speculate that the strong anti-correlation we found between individual salience for *Faces* and *Touched* reflects individual differences in developmental exposure and cortical layout. The same could apply to the anti-correlation between *Faces* and *Text*, two stimulus classes which have previously been proposed to compete for cortical territory^47^. It is worth noting that most of the reliable dimensions of individual salience differences we found correspond to domain specific patches of the ventral path (as is true for *Faces*^48–50^, *Text*^47,51^, *Motion*^52^, *Touched*^46,53,54^ and maybe *Taste*^55^). Our findings open up a whole new field of research, where the relationship between salience differences and the individual functional architecture of the ventral stream is directly compared.

Our findings raise important questions about the individual nature of visual perception. Two observers presented with the same image likely end up with a different percept^5,6^ and interpretation^56^ of this image when executing systematically different eye movements. Vision scientists may be chasing a phantom when ‘averaging out’ individual differences to study the ‘typical observer’^57–59^ and *vice versa* perception may be crucial to understand individual differences in cognitive abilities^60,61^, personality^62,63^, social behaviour^28,44^, clinical traits^64–66^ and development^45^.

We took a major first step towards uncovering the perceptual implications of salience differences here. Individual face salience was significantly correlated with face recognition skills. Interestingly, this was only true when considering the proportion of *first* fixations attracted by faces. Immediate saccades towards faces can have very short latencies and be under limited voluntary control^67,68^, likely reflecting bottom-up processing. This raises questions about the ontological interplay between face salience and recognition. Small initial differences may grow through mutual reinforcement of face fixations and superior perceptual processing.

To our surprise, we found no significant correlations between salience differences and personality factors captured by standard psychometric questionnaires. However, given the large number of tests and control for multiple testing, we cannot rule out the existence of small or medium effects. Either way, the salience differences we found should inform and enrich the science of individual differences. They have unusually high magnitudes (up to factor 3) and reliability (up to and greater than .9) for objective psychological traits and appear to document fundamental differences in visual attention and perception, which are main filters of incoming information about our surroundings

In summary, we found a small set of semantic dimensions that span a space of individual differences in fixation behaviour. These dimensions replicated across culturally diverse samples and also applied to the first fixations directed towards an image. Visual salience is not just a function of the image, but also of the individual observer.

## Methods

### Subjects

The study comprised three original datasets (the *Lon*, *Gi_1* and *Gi_2* samples) and the re-analysis of a public dataset (the *Xu et al*. sample^8^).

54 healthy adults with normal or corrected-to normal vision participated in the experiment conducted at University College London, UK. The local institutional review board approved the study and participants provided written informed consent. Reimbursement was 25£. Eyetracking failed for three participants due to technical problems, leaving 51 in the *Lon* sample (25 males, 4 left handed, mean age 23 with a standard deviation of 4 years). For two of these participants, data from one block was missing due to a recording failure; the remaining data from these participants were entered into the analysis nonetheless. Control-analysis showed that the results remained virtually unchanged when these data were excluded. 46 of the *Lon* participants additionally completed three personality questionnaires administered online: The IPIP-NEO 120 (NEO)^40^ to measure the Big Five^69^; the Highly Sensitive Person Scale (HSP)^42^ and the Brief Sensation Seeking Scale (BSSS)^41^.

51 healthy adults with normal or corrected-to normal vision participated in the experiment conducted at Justus-Liebig Universität Giessen, Germany (the *Gi_1* sample; 11 males, 5 left handed, mean age 24, with a standard deviation of 4 years). For one of these participants, data from one block was missing due to a recording failure; the remaining data from this participants were entered into the analysis nonetheless. Control-analysis showed that results remained virtually unchanged when these data were excluded. The local institutional review board approved the study and participants provided written informed consent. Reimbursement was 30€ or course credits.

48 of the *Gi_1* participants returned to the lab for a re-test (the *Gi_2* sample). The period between tests was 16 days on average (minimum 6 days; maximum: 43 days; standard deviation: 7 days). 46 of the *Gi* participants additionally completed an online version of the Cambridge Face Memory Test (CMFT)^70^.

The public *Xu et al*. dataset comprises 15 participants with a reported age range of 18-30 years^8^.

### Stimuli

Participants were presented with 700 natural images depicting a wide variety of complex everyday scenes (http://www-users.cs.umn.edu/~qzhao/predicting.html). Semantic metadata^8^ consist of binary pixel maps for 5551 objects in these images and accompanying labels for 12 semantic dimensions. We modified the provided labels to minimize overlap between them in the following way: The (neutral) *Faces* label was removed from all objects with the *Emotion* label (i.e. emotional faces); The *Smell* label was removed from all objects with the *Taste* label; The *Operable* and *Gazed* labels were removed from all objects with the *Touched* label; The *Watchable* label was removed from all objects with the *Text* label. This step allowed to test individual differences in fixation attraction for a given dimension largely independently from the others. Without this step some of the attributes would have been perfectly confounded (for instance, all text is watchable; all emotional faces are faces).

In each experiment, participants viewed seven blocks of 100 images on a screen. Stimulus presentation and data collection was coded in MATLAB Version R2016b (MathWorks, Natick, MA) using Psychtoolbox Version 3.0.12^71,72^. For the *Lon* and *Gi_1* samples the order of images was fixed across all participants and each image was presented for 3s, with a self-paced period of a central fixation dot in between. Participants were simply instructed to ‘look at the images in any way [they] want’ and initiated the onset of each image with a press of the space bar. The re-test of the Giessen sample (*Gi_2*) followed the same procedure as the first appointment, with the exception of the order of images presented, which was shuffled relative to the first appointment (but again constant across participants).

Participants in the *Lon* sample sat at a distance of ~ 65 cm from a screen and saw the stimuli at a resolution of 800 × 600 pixels and a size of 19.2 × 14.4 degrees visual angle. Participants in the *Gi* samples sat with their head in a chinrest, at a distance of 46 cm from the screen and saw the stimuli at a resolution and size of 1000×750 pixels and a size of 37.2 × 27.9 degrees visual angle. The *Xu et al*. sample saw stimuli at a size of 33.7 × 25.3 degrees visual angle. Further details on this sample can be found in the original publication by Xu et al.^8^.

### Data Collection

The gaze of participants in the *Lon* sample was sampled remotely and binocularly with a Tobii EyeX (Tobii Technologies, Danderyd, Sweden) at a frequency of ~ 55 Hz^73^. Eyetracking data from the *Gi* samples was collected from the left eye with a tower mounted Eyelink 1000 (SR Research, Ottawa, Canada) at a frequency of 2 kHz (the *Xu et al*. data were collected with a remote Eyelink 1000^8^).

At the beginning of each block, participants completed a nine-point calibration and validation procedure, which was repeated if necessary. For the *Gi* samples, fixation data was collected online using the ‘normal’ setting of the Eyelink parser (saccade velocity and acceleration thresholds of 30 d.v.a./s and 9500 d.v.a./s^2^, respectively) and the default drift check procedure in each inter-trial interval. Raw data from the *Lon* sample was converted to fixation data offline (see below).

### Data Processing and Analyses

Raw eyetracking data from the *Lon* sample were transformed into fixation data applying a saccade threshold of 30 d.v.a./s and taking the median x and y position of fixation samples as the respective fixation location. Fixations were drift-corrected in a block-wise fashion, centering on the median of fixation locations registered at image onsets during the respective block.

Onset fixations and fixations with a duration below 100ms were disregarded for all datasets (minimum fixation duration following standard recommendations by SR Research and Xu *et al*. ^8^). Fixations that fell on or within a distance of ~ 0.5 d.v.a from a labelled object were assigned the corresponding label. Unlabelled fixations were disregarded for the calculation of cumulative fixation times and the individual proportion of *first* fixations (see below).

In order to quantify the individual tendency to fixate objects bearing a given attribute label, we first calculated the cumulative fixation time for all labelled fixations made by a given observer to a given image set. This allowed us to calculate the proportion of this time spent on a given attribute in a second step.

The *first* fixations analysis considered the proportion of labelled *first* fixations (after image onset) landing on objects with a given attribute for a given observer and image set. Individual proportions of cumulative fixation time and first fixations were expressed in %.

### Consistency and re-test correlations

To estimate the consistency of individual differences in % cumulative fixation time or % first fixations along a given attribute dimension, we calculated these measures independently for two random halves of the image for each observer and then correlated individual differences for one half of the images with the other using Pearson correlation coefficients. This procedure was repeated 1000 times. We inspected the frequency histograms of all correlations (Fig. 2) and considered the median correlation coefficient across random image splits as an indicator of consistency. The accompanying two-sided *P*-values were Bonferroni adjusted for twelve comparisons in the *Lon* sample,

To estimate the re-test reliability of individual differences in % cumulative fixation time or % first fixations for a given attribute, we correlated the corresponding values for the first session of the *Gi* sample with those of the second.

### Covariance patterns between dimensions and correlations with other measures

To investigate the covariance pattern between the dimensions of consistent individual differences in fixation behaviour, we inspected pair-wise Pearson correlation matrices (Fig. 3B and 4, left hand side). We further used metric multidimensional scaling to project the pair-wise dissimilarities between dimensions (defined as (1-*r*)^2^) onto distances in a two-dimensional space (Fig. 3B and 4, right hand side). To further test the stability of covariance patterns between datasets we calculated pair-wise Pearson correlations between Fisher Z-transformed correlation matrices from different samples.

To explore the relationship between personality and individual gaze behaviour, we computed all pairwise correlations between reliable dimensions of salience differences and personality. Specifically we correlated individual salience differences for *Faces*, *Emotion*, *Text*, *Taste*, *Touched* and *Motion* with the Big Five as determined by the IPIP-NEO 120^40^ *Neuroticism* (N), *Extraversion* (E), *Openness* (O), *Assertiveness* (A), *Conscientiousness* (C), with high sensitivity (HS) as determined by the Highly Sensitive Person scale^42^, and with sensation seeking (SS) as determined by the Brief Sensation Seeking Scale^41^. The resulting 6×7 correlation matrix was computed separately for the proportion of cumulative fixation time and for that of *first* fixations; accompanying *p*-values were adjusted for multiple testing using the Holm-Bonferroni correction for family-wise errors^43^.

To test the perceptual implications of salience differences, we concentrated of the example of faces. Specifically, we probed a correlation between individual face salience as indicated by the proportion of cumulative dwell time or *first* fixations landing on faces and total scores in the Cambridge Face Memory Test^70^. This standard test of individual face recognition skills was administered online to participants from the *Gi* sample, after the first eyetracking session.

### Availability of data and code

Anonymised fixation data and code to reproduce the results presented here are freely available at https://osf.io/n5v7t/

## Acknowledgements

This work was supported by a JUST’US fellowship from the University of Giessen (B.d.H), as well as a research fellowship by the Deutsche Forschungsgemeinschaft (DFG HA 7574/1-1; B.d.H.). KRG was supported by DFG Collaborative Research Center SFB/TRR 135. We would like to thank Xu et al. for publishing their stimuli and dataset, Dr. Pete R. Jones for binding code bridging between the Eye-X C library and MATLAB, as well as Ms Diana Weissleder for help with collecting the *Gi* datasets.

## Author Contributions

Conception and design: BdH, Data Collection: ALI, BdH, Data Analysis: BdH, in discussion with KRG. Provision of lab space and resources: KRG, DSS. Manuscript writing: BdH wrote the initial draft. All authors discussed the results and final manuscript.

## Supplementary Information

### Supplementary Methods

To test the relationship between individual salience and visual field biases, we first explored the distribution of objects corresponding to dimensions of individual salience across image space (Supplementary Fig. 1a; neutral and emotional faces collapsed into *Faces*). The most prominent spatial bias in the scene stimuli was a strong tendency for faces to appear in the upper visual field (~ 75% of faces).

Therefore, we tested whether individual face salience (individual % of *first* fixations landing on faces) correlated with a general upper visual field bias. We defined the latter as the median elevation of *first* fixations towards images *not* containing a face (relative to central fixation and expressed in % image height). This was indeed the case (*r* = .49-.78 across samples, all *P* < .05; Supplementary Fig. 1b).

Given a correlation between individual face salience and a general bias towards the upper visual field, we wanted to test whether individual salience preferences *depended* on visual field biases. Therefore, we tested whether individual differences in face salience persisted for the lower visual field, correlating the individual percentage of first fixations landing on faces in the upper visual field (UVF) with that landing on faces in the lower visual field (LVF). For this analysis, we defined UVF and LVF faces as those for which the median height of the pixel mask was in the upper and lower image half, respectively. Importantly, individual face salience was stable across the UVF and LVF, showing that it persisted independently of the UVF bias (*r* = .61-.87 across samples, all *P* < .05; Supplementary Fig. 1c).

**Supplementary Figure 1.**
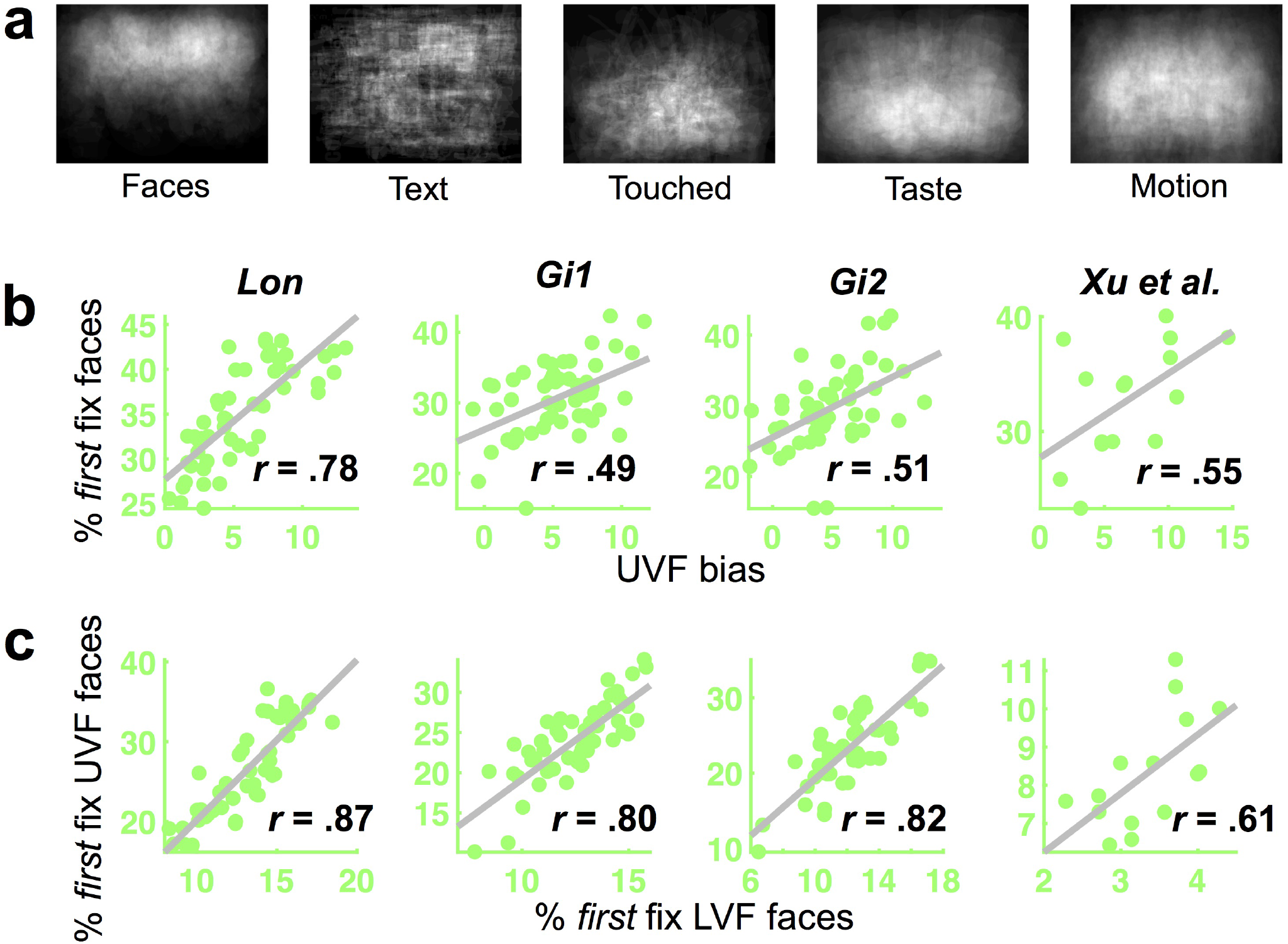
Individual salience and visual field biases. (a) Distribution of different types of objects in image space across scenes. Notice the upwards bias of Faces. **(b)** Individual face salience significantly correlated with a general upper visual field bias in all four samples (i.e. with the elevation of first fixations towards images without a face; see Suppelemtary Methods above for details). **(c)**. Nevertheless, individual face salience persisted independent of this spatial bias. The individual proportion of first fixations landing in the upper and lower visual field (UVF and LVF) were highly correlated with each other.

## References

1. Gegenfurtner, K. R. The Interaction Between Vision and Eye Movements. Perception 1–25 (2016). doi:10.1177/0301006616657097

2. Curcio, C. A. & Allen, K. A. Topography of ganglion cells in human retina. J. Comp. Neurol. 300, 525 (1990).

3. Dougherty, R. F. et al. Visual field representations and locations of visual areas V1/2/3 in human visual cortex. J. Vis. 3, 586–98 (2003).

4. Rosenholtz, R. Capabilities and Limitations of Peripheral Vision. Annu. Rev. Vis. Sci. 437–459 (2016). doi:10.1146/annurev-vision-082114-035733

5. Henderson, J. M., Williams, C. C., Castelhano, M. S. & Falk, R. J. Eye movements and picture processing during recognition. Percept. {&} Psychophys. 65, 725–734 (2003).

6. Nelson, W. W. & Loftus, G. R. The functional visual field during picture viewing. J. Exp. Psychol. Hum. Learn. 6, 391–399 (1980).

7. Harel, J., Koch, C. & Perona, P. Graph-Based Visual Saliency. Proceedings of the 19th International Conference on Neural Information Processing Systems 545–552 (2006).

8. Xu, J., Jiang, M., Wang, S., Kankanhalli, M. S. & Zhao, Q. Predicting human gaze beyond pixels. J. Vis. 14, (2014).

9. Henderson, J. M. & Hayes, T. R. Meaning-based guidance of attention in scenes as revealed by meaning maps. Nat. Hum. Behav. 1, 743–747 (2017).

10. Einhauser, W., Spain, M. & Perona, P. Objects predict fixations better than early saliency. J. Vis. 8, 18–18 (2008).

11. Borji, A. & Itti, L. Defending Yarbus: eye movements reveal observers’ task. J. Vis. 14, 29 (2014).

12. Tatler, B. W., Hayhoe, M. M., Land, M. F. & Ballard, D. H. Eye guidance in natural vision: Reinterpreting salience. J. Vis. 11, 5–5 (2011).

13. Itti, L., Koch, C. & Niebur, E. A model of saliency-based visual attention for rapid scene analysis. IEEE Trans. Pattern Anal. Mach. Intell. 20, 1254–1259 (1998).

14. White, B. J. et al. Superior colliculus neurons encode a visual saliency map during free viewing of natural dynamic video. Nat. Commun. 8, 14263 (2017).

15. Bogler, C., Bode, S. & Haynes, J.-D. Decoding successive computational stages of saliency processing. Curr. Biol. 21, 1667–71 (2011).

16. Stoll, J., Thrun, M., Nuthmann, A. & Einhäuser, W. Overt attention in natural scenes: objects dominate features. Vision Res. 107, 36–48 (2015).

17. Kummerer, M., Wallis, T. S. A., Gatys, L. A. & Bethge, M. Understanding Low- and High-Level Contributions to Fixation Prediction. in 2017 IEEE International Conference on Computer Vision (ICCV) 4799–4808 (IEEE, 2017). doi:10.1109/ICCV.2017.513

18. Kümmerer, M., Wallis, T. S. A. & Bethge, M. Information-theoretic model comparison unifies saliency metrics. Proc. Natl. Acad. Sci. U. S. A. 112, 16054–9 (2015).

19. Buswell, G. T. How people look at pictures: a study of the psychology and perception in art. How people look at pictures: a study of the psychology and perception in art. (Univ. Chicago Press, 1935).

20. Yarbus, A. L. in Eye Movements and Vision 171–211 (Springer US, 1967). doi:10.1007/978-1-4899-5379-7_8

21. Andrews, T. J. & Coppola, D. M. Idiosyncratic characteristics of saccadic eye movements when viewing different visual environments. Vision Res. 39, 2947–53 (1999).

22. Castelhano, M. S. & Henderson, J. M. Stable individual differences across images in human saccadic eye movements. Can. J. Exp. Psychol. 62, 1–14 (2008).

23. Henderson, J. M. & Luke, S. G. Stable individual differences in saccadic eye movements during reading, pseudoreading, scene viewing, and scene search. J. Exp. Psychol. Hum. Percept. Perform. 40, 1390–400 (2014).

24. Meyhöfer, I., Bertsch, K., Esser, M. & Ettinger, U. Variance in saccadic eye movements reflects stable traits. Psychophysiology 53, 566–578 (2016).

25. Rigas, I. & Komogortsev, O. V. Current research in eye movement biometrics: An analysis based on BioEye 2015 competition. Image Vis. Comput. (2016). doi:10.1016/j.imavis.2016.03.014

26. Bargary, G. et al. Individual differences in human eye movements: An oculomotor signature? Vision Res. 141, 157–169 (2017).

27. Ettinger, U. et al. Reliability of smooth pursuit, fixation, and saccadic eye movements. Psychophysiology 40, 620–628 (2003).

28. Constantino, J. N. et al. Infant viewing of social scenes is under genetic control and is atypical in autism. Nature 547, 340–344 (2017).

29. Kennedy, D. P. et al. Genetic Influence on Eye Movements to Complex Scenes at Short Timescales. Curr. Biol. 27, 3554–3560.e3 (2017).

30. Parkhurst, D., Law, K. & Niebur, E. Modeling the role of salience in the allocation of overt visual attention. Vision Res. 42, 107–23 (2002).

31. Foulsham, T. & Underwood, G. What can saliency models predict about eye movements? Spatial and sequential aspects of fixations during encoding and recognition. J. Vis. 8, 6 (2008).

32. Einhauser, W., Rutishauser, U. & Koch, C. Task-demands can immediately reverse the effects of sensory-driven saliency in complex visual stimuli. J. Vis. 8, 2 (2008).

33. Anderson, N. C., Ort, E., Kruijne, W., Meeter, M. & Donk, M. It depends on *when* you look at it: Salience influences eye movements in natural scene viewing and search early in time. J. Vis. 15, 9 (2015).

34. Mackay, M., Cerf, M. & Koch, C. Evidence for two distinct mechanisms directing gaze in natural scenes. J. Vis. 12, 9–9 (2012).

35. Henderson, J. M. & Hayes, T. R. Meaning guides attention in real-world scene images: Evidence from eye movements and meaning maps. J. Vis. 18, 10 (2018).

36. Nyström, M. & Holmqvist, K. Semantic Override of Low-level Features in Image Viewing – Both Initially and Overall. J. Eye Mov. Res. 2, (2008).

37. Kaiser, D. & Cichy, R. M. Typical visual-field locations enhance processing in object-selective channels of human occipital cortex. J. Neurophysiol. 120, 848–853 (2018).

38. de Haas, B. et al. Perception and Processing of Faces in the Human Brain Is Tuned to Typical Feature Locations. J. Neurosci. 36, (2016).

39. de Haas, B. & Schwarzkopf, D. S. Feature-location effects in the Thatcher illusion. J. Vis. 18, (2018).

40. Maples, J. L., Guan, L., Carter, N. T. & Miller, J. D. A test of the International Personality Item Pool representation of the Revised NEO Personality Inventory and development of a 120-item IPIP-based measure of the five-factor model. Psychol. Assess. 26, 1070–84 (2014).

41. Hoyle, R. H., Stephenson, M. T., Palmgreen, P., Lorch, E. P. & Donohew, R. L. Reliability and validity of a brief measure of sensation seeking. Pers. Individ. Dif. 32, 401–414 (2002).

42. Aron, E. N. & Aron, A. Sensory-processing sensitivity and its relation to introversion and emotionality. J. Pers. Soc. Psychol. 73, 345–368 (1997).

43. Holm, S. A simple sequentially rejective multiple test procedure. Scand. J. Stat. (1979).

44. Wang, S. et al. Atypical Visual Saliency in Autism Spectrum Disorder Quantified through Model-Based Eye Tracking. Neuron 88, 604–16 (2015).

45. Jones, W. & Klin, A. Attention to eyes is present but in decline in 2-6-month-old infants later diagnosed with autism. Nature 504, 427–31 (2013).

46. Arcaro, M. J., Schade, P. F., Vincent, J. L., Ponce, C. R. & Livingstone, M. S. Seeing faces is necessary for face-domain formation. Nat. Neurosci. 20, 1404–1412 (2017).

47. Dehaene, S. et al. How learning to read changes the cortical networks for vision and language. Science 330, 1359–64 (2010).

48. Kanwisher, N. & Yovel, G. The fusiform face area: a cortical region specialized for the perception of faces. Philos. Trans. R. Soc. Lond. B. Biol. Sci. 361, 2109–28 (2006).

49. Tsao, D. Y., Moeller, S. & Freiwald, W. A. Comparing face patch systems in macaques and humans. Proc. Natl. Acad. Sci. U. S. A. 105, 19514–9 (2008).

50. Grill-Spector, K. & Weiner, K. S. The functional architecture of the ventral temporal cortex and its role in categorization. Nat. Rev. Neurosci. 15, 536–548 (2014).

51. McCandliss, B. D., Cohen, L. & Dehaene, S. The visual word form area: expertise for reading in the fusiform gyrus. Trends Cogn. Sci. 7, 293–299 (2003).

52. Kourtzi, Z. & Kanwisher, N. Activation in human MT/MST by static images with implied motion. J. Cogn. Neurosci. 12, 48–55 (2000).

53. Orlov, T., Makin, T. R. & Zohary, E. Topographic representation of the human body in the occipitotemporal cortex. Neuron 68, 586–600 (2010).

54. Weiner, K. S. & Grill-Spector, K. Neural representations of faces and limbs neighbor in human high-level visual cortex: evidence for a new organization principle. Psychol. Res. 77, 74–97 (2013).

55. Adamson, K. & Troiani, V. Distinct and overlapping fusiform activation to faces and food. Neuroimage 174, 393–406 (2018).

56. Bush, J. C., Pantelis, P. C., Morin Duchesne, X., Kagemann, S. A. & Kennedy, D. P. Viewing Complex, Dynamic Scenes “Through the Eyes” of Another Person: The Gaze-Replay Paradigm. PLoS One 10, e0134347 (2015).

57. Wilmer, J. B. How to use individual differences to isolate functional organization, biology, and utility of visual functions; with illustrative proposals for stereopsis. Spat. Vis. 21, 561–79 (2008).

58. Charest, I. & Kriegeskorte, N. The brain of the beholder: honouring individual representational idiosyncrasies. Lang. Cogn. Neurosci. 30, 367–379 (2015).

59. Peterzell, D. Discovering Sensory Processes Using Individual Differences: A Review and Factor Analytic Manifesto. Electron. Imaging 1–11 (2016).

60. Haldemann, J., Stauffer, C., Troche, S. & Rammsayer, T. Processing Visual Temporal Information and Its Relationship to Psychometric Intelligence. J. Individ. Differ. 32, 181–188 (2011).

61. Hayes, T. R. & Henderson, J. M. Scan patterns during real-world scene viewing predict individual differences in cognitive capacity. J. Vis. 17, 23 (2017).

62. Troche, S. J. & Rammsayer, T. H. Attentional blink and impulsiveness: evidence for higher functional impulsivity in non-blinkers compared to blinkers. Cogn. Process. 14, 273–81 (2013).

63. Wu, D. W.-L., Bischof, W. F., Anderson, N. C., Jakobsen, T. & Kingstone, A. The influence of personality on social attention. Pers. Individ. Dif. 60, 25–29 (2014).

64. Hayes, T. R. & Henderson, J. M. Scan patterns during scene viewing predict individual differences in clinical traits in a normative sample. PLoS One 13, e0196654 (2018).

65. Armstrong, T. & Olatunji, B. O. Eye tracking of attention in the affective disorders: a meta-analytic review and synthesis. Clin. Psychol. Rev. 32, 704–23 (2012).

66. Molitor, R. J., Ko, P. C. & Ally, B. A. Eye Movements in Alzheimer’s Disease. J. Alzheimers. Dis. 44, 1 (2015).

67. Crouzet, S. M., Kirchner, H. & Thorpe, S. J. Fast saccades toward faces: Face detection in just 100 ms. J. Vis. 10, 1–17 (2010).

68. Rösler, L., End, A. & Gamer, M. Orienting towards social features in naturalistic scenes is reflexive. PLoS One 12, e0182037 (2017).

69. Costa, P. T. & McCrae, R. R. Four ways five factors are basic. Pers. Individ. Dif. 13, 653–665 (1992).

70. Duchaine, B. & Nakayama, K. The Cambridge Face Memory Test: results for neurologically intact individuals and an investigation of its validity using inverted face stimuli and prosopagnosic participants. Neuropsychologia 44, 576–85 (2006).

71. Kleiner, M., Brainard, D. & Pelli, D. What’s new in Psychtoolbox-3? Perception 36, 14 (2007).

72. Pelli, D. G. The VideoToolbox software for visual psychophysics: transforming numbers into movies. Spat. Vis. 10, 437–42 (1997).

73. Gibaldi, A., Vanegas, M., Bex, P. J. & Maiello, G. Evaluation of the Tobii EyeX Eye tracking controller and Matlab toolkit for research. Behav. Res. Methods 49, 923–946 (2017).

